# Deciphering the Interaction Between Dvl2 and Profilin2: Effector Molecules of Non-Canonical Wnt Signaling

**DOI:** 10.1101/2025.10.09.681388

**Authors:** Saikat Das, Shubham Das, Sankar Maiti

## Abstract

Wnt signalling is a cornerstone of embryonic development, orchestrating critical processes such as body axis formation, gastrulation, and organogenesis through conserved canonical and non-canonical pathways. Dishevelled (Dvl), a central mediator of these pathways, contains conserved DIX, PDZ, and DEP domains, along with an extreme-C-terminus. Recent studies suggest that the extreme-C-terminus regulates non-canonical Wnt signalling via an autoinhibitory interaction with the PDZ domain. Non-canonical Wnt signalling branches into the planar cell polarity (PCP) and Wnt/Ca²⁺ pathways. Profilin, a monomeric actin-binding protein, has been implicated in PCP signalling through Daam1-mediated actin polymerization, whereas its silencing disrupts the Wnt/Ca²⁺ pathway in a Daam1-independent manner, pointing to a role for profilin upstream of Daam1. In this study, we identify a novel interaction between Dvl2 and profilin2. Co-localization and in vitro pull-down assays demonstrate that profilin2 directly interacts with Dvl2. Furthermore, our study reveals profilin2 binds specifically to the extreme-C-terminus of Dvl2, beyond the polyproline motif, without engaging the PDZ or DEP domains. This challenges the conventional view of profilin-polyproline interactions and highlights the existence of previously unrecognized molecular determinants. Moreover, we show that Dvl2 adopts an autoinhibited conformation through intramolecular binding of its extreme-C-terminus to the PDZ domain. Remarkably, profilin2 retains its binding ability even in this autoinhibited state. Together, these findings uncover a previously unrecognized profilin2-Dvl2 interaction and provide new mechanistic insights into the molecular regulation of non-canonical Wnt signalling.

## 1. Introduction

Embryonic development relies on precise, timely, and specific cell-to-cell communication. Molecular genetic studies have identified a set of conserved cores signalling pathways, including Wnt, Notch, Transforming Growth Factor beta (TGF-β), and Sonic-Hedgehog (Shh) [1–4]. Wnt signalling pathway is highly conserved across metazoan animals, plays a critical role in body axis formation, gastrulation, germ layer specification, and early organogenesis [5,6]. Wnt signalling pathway is broadly classified into two groups, canonical and non-canonical Wnt signalling [7,8]. The canonical pathway governs cell fate and contributes to dorsal axis formation, while the non-canonical pathway regulates convergent extension during gastrulation and tissue mophogenesis [9,10]. Wnt signalling is modulated by various regulators categorized as extracellular, membrane, and cytoplasmic. Extracellular regulators like Wnt ligands and R-spondins act as agonists, promoting signalling, while Dickkopf (DKK) and Secreted Frizzled-related proteins (sFRPs) serve as antagonists, inhibiting Wnt activity [11–13]. At the membrane, membrane regulators like Frizzled (Fz) receptors and co-receptors such as LRP5 and ROR regulate signal transduction [14]. Dishevelled (Dvl) is a key cytoplasmic regulator facilitating Wnt pathway activation based on specific Wnt signals [15]. Notably, depletion of Dvl in Xenopus embryos results in disruption of gastrulation due to inhibition of convergent extension [16]. Moreover, there are three Dvl isoforms (Dvl1, Dvl2, and Dvl3) in humans [17]. Among the three isoforms Dvl2 is the most abundantly expressed [18]. Dvl1 and Dvl2 are directly involved in regulating Wnt signalling, whereas the role of Dvl3 is less well-defined [19]. Dvl2 contains conserved DIX (Dishevelled/Axin), PDZ (PSD-95, DLG, ZO1), and DEP (Dishevelled, EGL-10, Pleckstrin) domains [20]. The DIX domain facilitates canonical Wnt signalling by preventing the β-catenin destruction complex from targeting β-catenin for proteasomal mediated degradation [21]. The accumulated β-catenin moves to nucleus where it activates target genes [22].

Non-canonical Wnt signalling consists of two distinct pathways, the planar cell polarity (PCP) pathway and the Wnt/Ca^2+^ pathway [23]. PCP signalling is initiated by forming asymmetric distribution of proteins within a cell. Distally, Fz, Dvl, and Diego (Dgo, or Diversin in vertebrates) form a complex, while proximally, the complex of Strabismus (Stbm, or Vangl in vertebrates) and Prickle (Pk) are localized [24]. Upon binding of Wnt ligand to Fz, Dvl2 forms a complex with Dishevelled-associated activator of morphogenesis 1 (Daam1), leading to Daam1 activation and subsequent RhoA activation via a Rho guanine exchange factor [25]. Daam1, a formin family protein, contains two conserved formin homology domains, FH1 and FH2 [26]. Formins act as nucleation promoting factors (NPFs), accelerating actin polymerization [27]. The FH1 domain, rich in polyproline, interacts with profilin, a monomeric actin-binding protein, while the FH2 domain binds actin to promote polymerization [28]. Activated RhoA triggers Rho-associated kinase (ROCK), which drives actin cytoskeletal reorganization [25]. Whereas, in the Wnt/Ca^2+^ signaling pathway, Dvl2, through trimeric G-protein, activates phospholipase C (PLC), which cleaves phosphatidylinositol 4,5-bisphosphate (PIP_2_) into inositol-1,4,5-trisphosphate (IP_3_) and diacylglycerol (DAG) [29]. IP_3_ induces the release of intracellular Ca^2+^ ions, which activate calcium/calmodulin-dependent kinase II (CamKII), TGF-β-activated kinase (TAK), and Nemo-like kinase (NLK), ultimately suppressing β-catenin-dependent gene expression [30]. Additionally, the elevated Ca^2+^ levels activate Protein kinase C (PKC) and Cdc42, promoting cytoskeletal reorganization [31].

Although the PDZ and DEP domains are essential for non-canonical Wnt signalling [16], the role of the C-terminal region following the DEP domain in Dvl on non-canonical Wnt signalling remain unexplored. Qi *et al.,* revealed that the C-terminus of mouse Dvl1 plays a role in regulation of non-canonical Wnt signalling [32]. Specifically, the extreme-C-terminus of mouse Dvl1 contains a class III PDZ-binding motif, which enables an autoinhibited state through intramolecular interaction with its own PDZ domain [32]. When an extreme-C-terminus deleted construct was overexpressed in a zebrafish embryo, canonical Wnt signalling was suppressed, and strong convergent extension (CE) defects were observed, indicating disruption of non-canonical Wnt signalling [33]. This highlights the essential function of the extreme-C-terminus in controlling non-canonical Wnt signalling. It also sparks questions about the possible role of other proteins might be involved in the above complex.

Profilin, a structurally conserved protein with four mammalian isoforms (Profilin1, Profilin2, Profilin3, Profilin4), plays a critical role in multiple developmental processes by binding various ligands [34]. It facilitates the exchange of ADP for ATP in monomeric actin [35]. Profilin also interacts with the FH1 domain of formins (like Daam1), supplying monomeric actin to the FH2 domain to support actin elongation [36]. Genetic studies in Drosophila reveal profilin’s essential roles in oogenesis, spermatogenesis, and bristle and eye formation [37,38]. Furthermore, profilin1 is vital for cytokinesis, with its deletion leading to embryonic lethality in mice at the two- to eight-cell stage, underscoring its importance in early development [39]. Profilin’s role in Wnt signalling was first observed in Xenopus embryo. Where it was shown that profilin2 was able to interact with the FH1 domain of Daam1 [40]. Overexpression of profilin2 in dorsal blastomeres of Xenopus embryos disrupts neural fold closure, while its downregulation in Keller’s explants inhibits convergent extension [40]. This hints towards profilin’s functions as a downstream effector of Daam1 in PCP pathway. Conversely, another study demonstrates silencing profilin1 in the T24M cell line suppresses Wnt/Ca^2+^ signalling by inhibiting PLC [41], suggesting profilin functions upstream of Daam1 along the Dvl axis. Therefore, we raise the question of whether profilin interacts with Dvl2.

## 2. Result

### 2.1. Dvl2 and Profilin2 Exhibit Subcellular Colocalization

To investigate the association between Dvl2 and profilin2, we generated antibodies against Dvl2 in mice and profilin2 in rabbits. Using terminal bleed, immunoblot analysis of purified proteins verified the specificity of both antibodies (Figure S1A & C). We further validated the functionality of the Dvl2 antibody, as our immunoblot of HEK293T cell lysates revealed a distinct Dvl2 band at approximately 90 kDa (Figure S1B).

To assess the co-localization of Dvl2 and profilin2, we examined their endogenous distribution in HEK293T cells using immunofluorescence staining followed by confocal microscopy. Dvl2 was detected in both the cytoplasm and nucleus, aligning with earlier reports (Figure 1A) [42,43]. Notably, unlike in cells overexpressing Dvl2, we did not observe any punctate cytosolic structures [42]. Moreover, profilin2 showed a distribution across both the nucleus and cytoplasm of the cell, in line with prior observations (Figure 1A) [44]. Further analysis of composite images (Dvl2 and Profilin2) revealed strong co-localization, particularly evident in the perinuclear region, as highlighted in the enlarged inset view (Figure 1A). This co-localization suggests profilin2’s role in functional activity of Dvl2.

**Figure 1.**
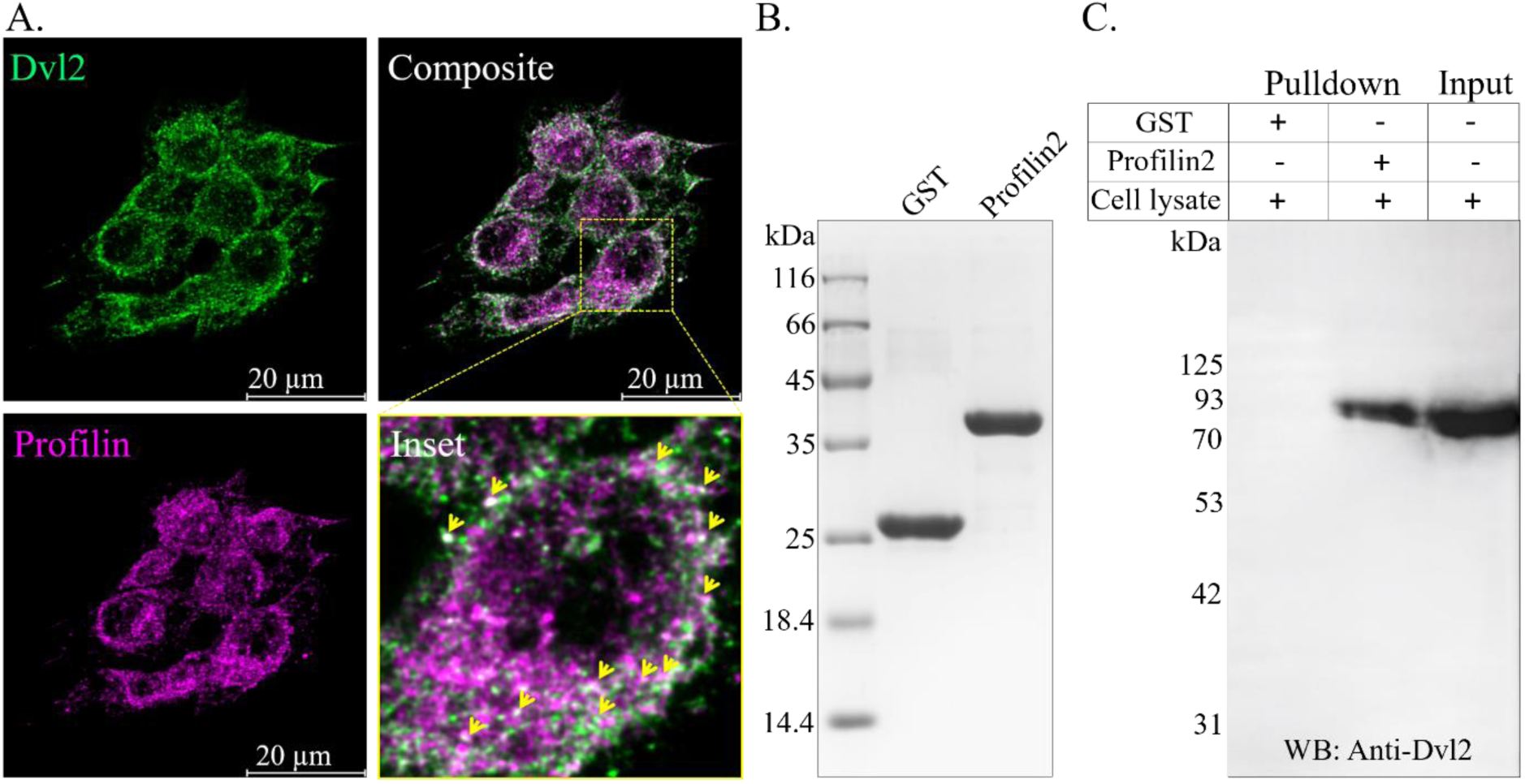
Dvl2 directly interacts Profilin2. (A) Immunofluorescence staining of Dvl2 and profilin2 in HEK293T cells was performed by fixing cells using a 1:1 ice-cold acetone-methanol mixture. Dvl2 and profilin2 were visualized with anti-Dvl2 and anti-profilin2 antibodies, respectively, while nuclei were stained with DAPI. Co-localization of Dvl2 and profilin2, indicated by arrows in the inset. The image represents a single Z-stack section. Scale bar = 20 µm. (B) Coomassie stained 10% gel showing the purified GST and GST-tagged profilin2. (C) Profilin2 interacts with Dvl2 in GST pull down assay. GST (negative control) and GST-profilin2 served as baits to capture Dvl2 (prey) from HEK293T cell lysate. Probing of Dvl2 in the samples was performed by western blot using anti-Dvl2 antibodies. GST-pull down samples are presented on the left while input samples are shown on the right.

### 2.2. Dvl2 Directly Binds to Profilin2

Our co-localization study demonstrated Dvl2 co-localized to profilin2. To observe the complex of Dvl2 and profilin2 in vitro, we conducted a in vitro pull-down assay. We purified GST-tagged profilin2 and GST alone (Figure 1B), utilizing them as bait and negative control, respectively (Figure 1C). Immunoblot analysis of the pull-down samples revealed that Dvl2 co-precipitated with GST-tagged profilin2, while no Dvl2 was detected with GST alone (Figure 1C). The input samples confirmed the presence of Dvl2 in the lysate (Figure 1C). These findings indicate a direct interaction between Dvl2 and profilin2.

### 2.3. C-terminal Fragment of Dvl2 Mediates Its Interaction with Profilin2

Prior studies have implicated the role of profilin2 in non-canonical Wnt signalling pathways [40]. The PDZ and DEP domains of Dvl2 have also been implicated in non-canonical Wnt signalling [45]. Our findings indicate a direct interaction between Dvl2 and profilin2. Therefore, to identify the region of Dvl2 involved in its interaction with profilin2, ELISA-based binding assays were performed following previously published protocol with slight modifications [46]. Based on domain organization, three Dvl2 constructs were generated in the pET28a+ vector. These includes Dvl2 (amino acids 182-389), which encompasses the PDZ domain, Dvl2 (amino acids 390-536), which contains the DEP domain, and Dvl2 (amino acids 536-736), representing the C-terminal region (Figure 2A). All constructs were expressed in *E. coli* as 6x-His-tagged proteins and purified. The molecular weights of Dvl2 (amino acids 182-389), Dvl2 (amino acids 390-536), and Dvl2 (amino acids 536-736), were approximately 25.7 kDa, 19 kDa, and 25 kDa, respectively (Figure 2B).

**Figure 2.**
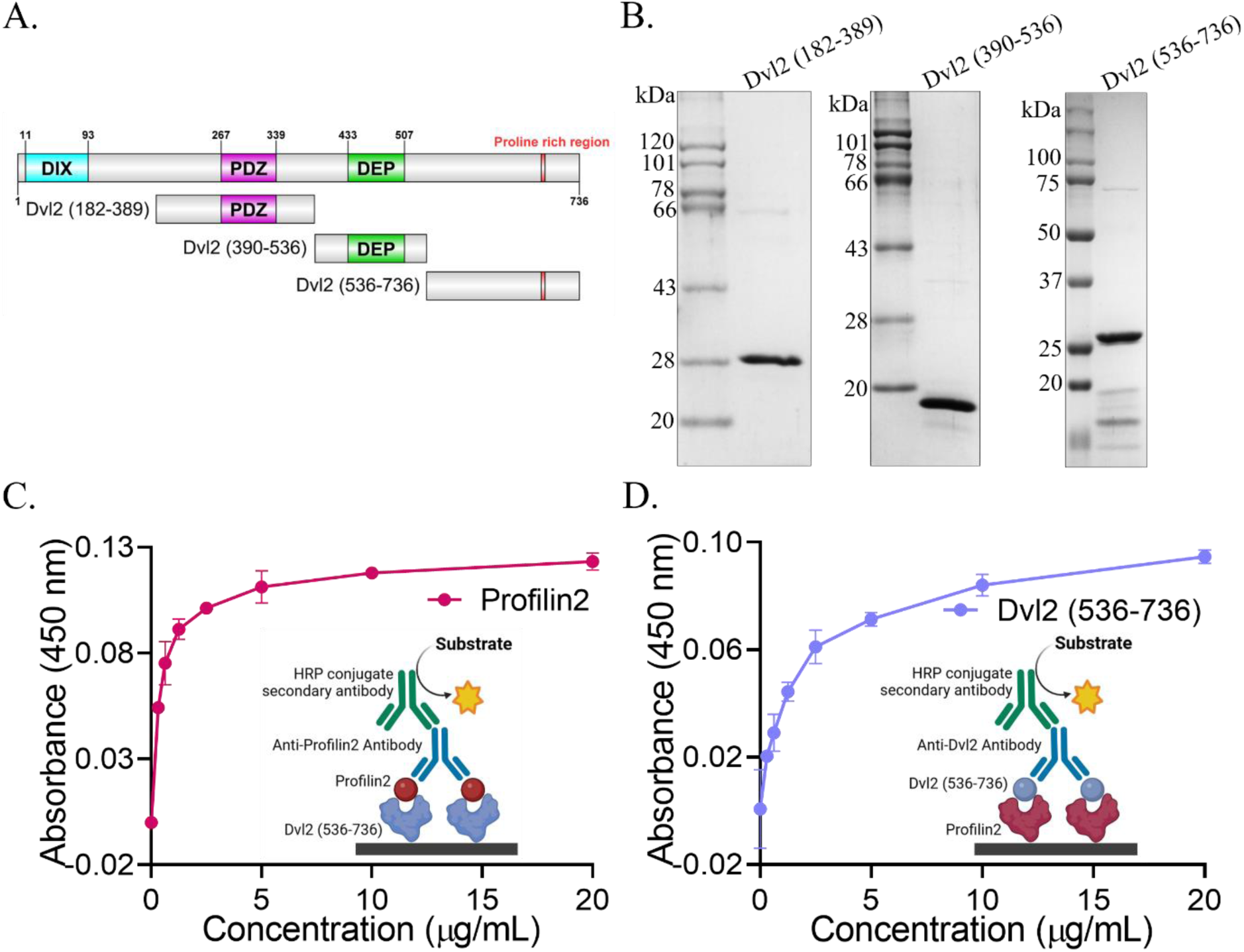
C-terminal Dvl2 (536-736) interacts with Profilin2. (A) Schematic representation of Dvl2 constructs. (B) Coomassie-stained 10% SDS-PAGE showing purified 6×His-tagged Dvl2 fragments: Dvl2 (amino acids 182-389), Dvl2 (amino acids 390-536), and Dvl2 (amino acids 536-736). (C) ELISA results for the interaction between Dvl2 (amino acids 536-736) and profilin2 where Dvl2 (amino acids 536-736) was coated as the ligand, with profilin2 added as the analyte in serial dilution. Rabbit anti-profilin2 antibody (1:1000) was used, followed by HRP-conjugated anti-IgG (1:10,000). (D) In the reverse setup, profilin2 was coated, and Dvl2 (amino acids 536-736) was added as the analyte. Mouse anti-Dvl2 antibody (1:500) was used, followed by HRP-conjugated anti-IgG (1:50,000). In both experiments, TMB was used as the substrate after adding HRP-conjugated anti-IgG, and absorbance was measured at 450 nm. Absorbance values were plotted on the Y-axis, and analyte concentrations were plotted on the X-axis.

Initially, we coated Dvl2 (amino acids 536-736) onto 96-well ELISA plates and introduced profilin2 as the analyte. We observed a concentration-dependent rise in absorbance that reached saturation, signifying the formation of a complex between this Dvl2 (amino acids 536-736) and profilin2 (Figure 2C). Subsequently, purified Dvl2 (amino acids 182-389) and Dvl2 (amino acids 390-536) fragments were coated and profilin2 was added as analyte (Figure S2). No notable absorbance was recorded, indicating that profilin2 does not form complex with the PDZ and DEP domain containing fragments Dvl2 (amino acids 182-389) and Dvl2 (amino acids 390-536) (Figure S2). These findings imply that profilin2 selectively binds to the C-terminal Dvl2 (amino acids 536-736).

To further validate this interaction, reciprocal ELISA experiments were performed using the Dvl2 (amino acids 536-736) fragment and GST-profilin2. In the first setup, Dvl2 (amino acids 536-736) was used as the ligand coated onto the wells, and profilin2 was added as the analyte (Figure 2C). In the second setup, the roles were reversed: profilin2 was coated onto the wells and Dvl2 (amino acids 536-736) served as the analyte (Figure 2D). In both configurations, a concentration-dependent increase in absorbance with saturation was observed. This indicates that, in the first setup, profilin2 forms a complex with Dvl2 (amino acids 536-736) (Figure 2C), and in the second setup, Dvl2 (amino acids 536-736) forms a complex with profilin2 (Figure 2D). These results collectively confirm that profilin2 interacts with C-terminal region of Dvl2, and PDZ, DEP domains have no role in the interaction with profilin.

### 2.4. Profilin2 interacts with the extreme-C-terminus of Dvl2 Beyond the Proline-rich Domain

The interaction between Profilin and polyproline-rich regions is well-documented. Enabled/vasodilator-stimulated phosphoprotein (Ena/VASP) is a well-known ligand for Profilin. The central proline-rich domain of VASP includes three GP5 motifs (GPPPPP), which bind directly to Profilin, increasing its affinity for actin and promoting filament elongation [47]. Microinjection of synthetic GP5 peptides into lateral amygdala (LA) neurons in mice disrupts the Ena/VASP-Profilin interaction, leading to impaired long-term memory formation [48]. This highlights the importance of profilin-polyproline region interaction. Our previous data demonstrated that profilin2 interacts with the C-terminal Dvl2 (amino acids 536-736), which contains a polyproline region (amino acids 686-691). This raises the question of whether the polyproline region plays a critical role in facilitating the interaction. To address this, we performed molecular docking between profilin2 and the C-terminal Dvl2 (amino acids 536-736). For that reason, we predicted 3D structure of Dvl2 (amino acids 536-736) using AlphaFold Colab, while the crystal structure of profilin2 was retrieved from the Protein Data Bank (PDB ID-2V8C). The findings indicated that profilin2 forms a complex with the C-terminal Dvl2 (amino acids 536-736) (Figure 3A). For further analysis, we chose the best model in an unbiased manner. Analysis of the best complex revealed a predominantly hydrophobic binding interface with five hydrogen bonds. The key interacting residues forming hydrogen bonds were Pro694, Ala695, Val696, Pro728, and Ser729 from Dvl2, and Gln4, Gln40, Asp48, Lys53, and Tyr78 from profilin2 (Figure 3A & S3). Notably, the region of Dvl2 that interacts with profilin2 lies beyond the polyproline region and includes residues at the extreme-C-terminus.

**Figure 3.**
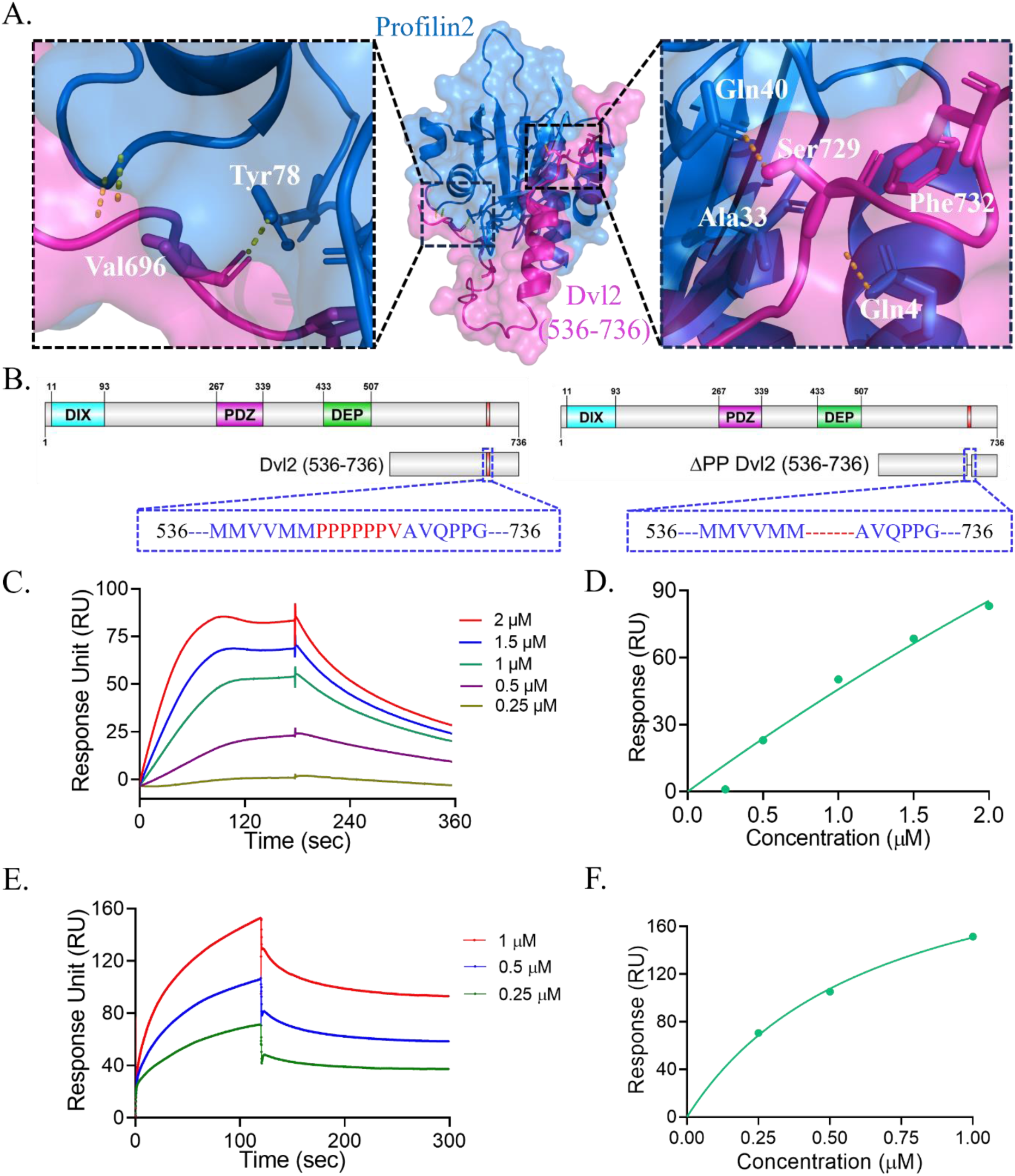
Profilin2 interacts with the extreme-C-terminus of Dvl2 in non-classical binding mode. (A) The best docked model of the Dvl2 (amino acids 536-736) (in magenta) and profilin2 (in marine blue) complex, with insets (left and right) highlighting the binding interface and interacting amino acids. (B) Diagram illustrating the domain organization of full-length Dvl2, Dvl2 (amino acids 536-736), and ΔPP C-terminal Dvl2 (amino acids 536-736) constructs, with a magnified view of the polyproline region (in red), emphasizing key amino acids within a blue dashed line box. SPR sensorgrams showing binding kinetics between Dvl2 and profilin2. Profilin2 was immobilized on a CM5 chip, while (C) Dvl2 (amino acids 536-736) and (E) ΔPP C-terminal Dvl2 (amino acids 536-736) served as analytes. The sensorgrams display concentration-dependent interactions of Dvl2 (amino acids 536-736) (C) and ΔPP C-terminal Dvl2 (amino acids 536-736) (E) with profilin2. The sensorgrams were fitted using a 1:1 Langmuir binding model. (D) & (F) Binding affinity curves of (C) and (E) respectively, where the X-axis showing concentration (µM) and the Y-axis showing response (RU). Curve fitting was conducted using a non-linear regression equation (one-site specific binding model).

To further validate our findings, we created a deletion construct, ΔPP C-terminal Dvl2 (amino acids 536-736), in which the polyproline region spanning amino acids 686-691 was deleted (Figure 3B). We then assessed the interaction between profilin2 and both the wild-type and deletion (ΔPP C-terminal Dvl2 (amino acids 536-736)) constructs using surface plasmon resonance (SPR) with previously published protocol [49]. Profilin2 was immobilized on a CM5 sensor chip (Figure S4A & B), and purified Dvl2 (amino acids 536-736) and ΔPP C-terminal Dvl2 (amino acids 536-736) (Figure S4C) were used as analytes. A significant binding response was observed in both cases. The kinetic parameters for the wild-type Dvl2 (amino acids 536-736) were, K_D_ = 217 nM, k_a_ = 0.512 × 10⁵ M⁻¹s⁻¹, k_d_ = 1.067 × 10⁻³ s⁻¹ (Figure 3C & D). For the polyproline-deleted construct, the parameters were, K_D_ = 255 nM, k_a_ = 0.107 × 10⁵ M⁻¹s⁻¹, k_d_ = 2.736 × 10⁻³ s⁻¹ (Figure 3E & F). These results clearly indicate that the deletion of the polyproline region does not intervene the interaction of profilin2 and Dvl2.

In corroboration with both datasets, we demonstrate for the first time that Profilin preferentially interacts with a region outside the classical polyproline motif, despite its presence of a polyproline region, suggesting a novel mode of Dvl2-Profilin interaction. This indicates profilin2 appears to favour an alternative binding surface for interaction with Dvl2.

### 2.5. Profilin2 Is Incorporated into the Autoinhibitory Architecture of Dvl2

Dvl2 is a key regulator of both canonical and non-canonical Wnt signalling pathways [50]. Dvl2 contains three conserved domains, DIX, PDZ, and DEP and its extreme-C-terminus shares homology across species [45]. To assess the conservation of the extreme-C-terminus of human Dvl2, we first performed a multiple sequence alignment of full-length Dishevelled isoforms from human (Dvl1, Dvl2, Dvl3), mouse (Dvl1, Dvl2, Dvl3), Xenopus (Dvl1, Dvl2, Dvl3), zebrafish (Dvl1, Dvl2, Dvl3), drosophila (Dsh), ascidian (Dvl), and Hydra (Dsh) (Figure S5). The analysis revealed a high degree of conservation in the DIX, PDZ, and DEP domains of human Dvl2 (Figure S5). Interestingly, the alignment also indicated notable conservation at within the last 40 amino acids of Dishevelled, which are part of the extreme-C-terminus.

To further investigate the conservation of this region, we conducted a separate multiple sequence alignment focusing on the last 40 amino acids of Dishevelled from the aforementioned species. Results showed that the extreme-C-terminus of human Dvl2 shares over 80% sequence similarity with its counterparts in mouse, Xenopus, and zebrafish, and over 60% similarity with those from ascidian and Hydra. Except, this region is poorly conserved in Drosophila (Figure 4A). This drastic sequence variation prompted us to investigate further, focusing on the extreme-C-terminus of Drosophila Dsh.

**Figure 4.**
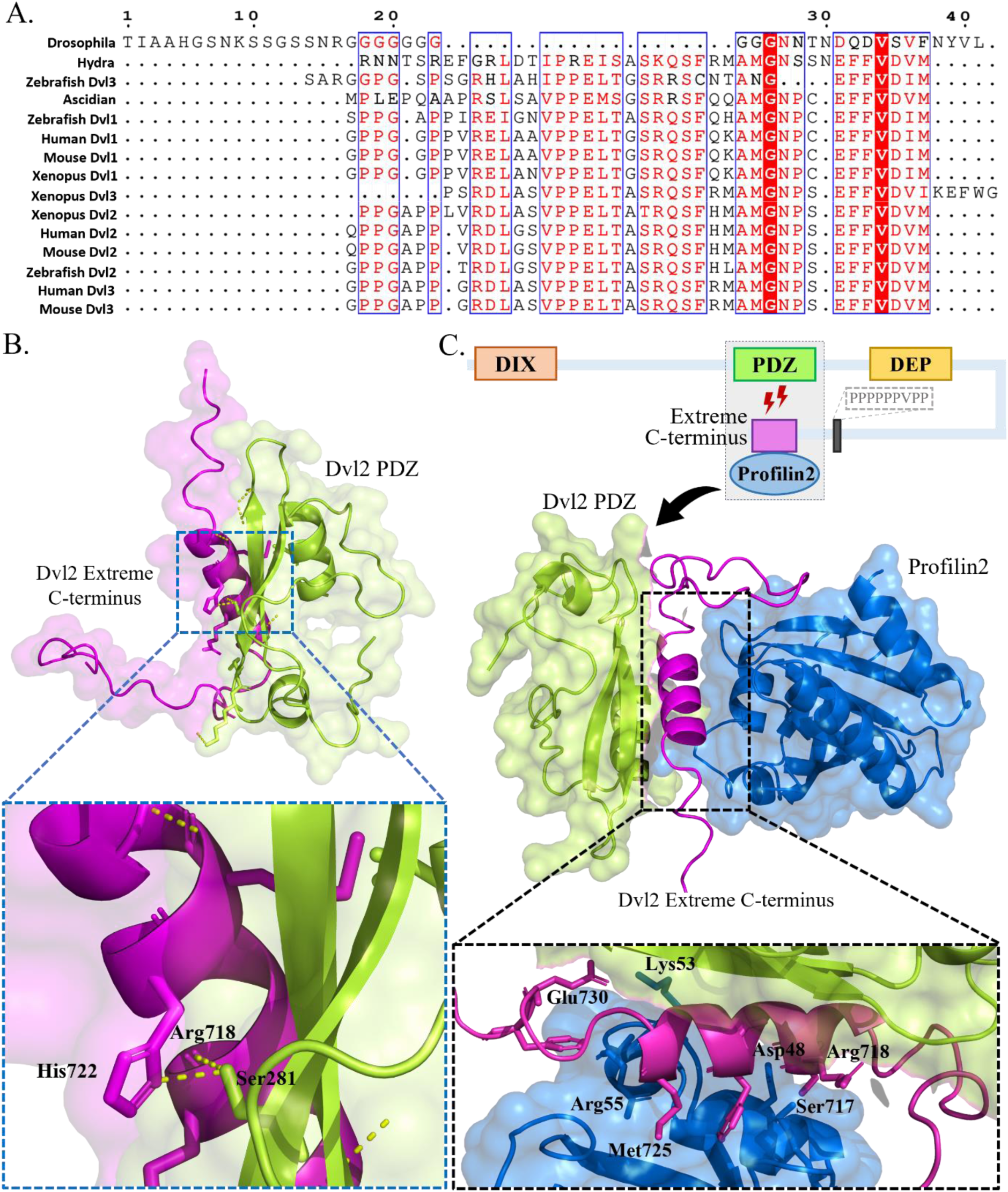
Dvl2 PDZ and extreme-C-terminus interaction remains unaffected by Profilin2 binding. (A) Multiple sequence alignment of the last 40 amino acids of Dishevelled from Hydra, ascidian, Drosophila, zebrafish, Xenopus, mouse, and human. Residues conserved across animal classes are shown as red letters within blue boxes, while fully conserved residues appear as white letters on a red background. (B) The best view of the autoinhibitory complex between the Dvl2 PDZ domain (in lime green) and its extreme-C-terminus (in magenta), with a zoomed-in view highlighting key amino acid interactions at the binding interface. (C) Schematic illustrating profilin2’s involvement in the PDZ-extreme-C-terminus complex. The best docking pose of profilin2 (in marine blue) shows that its association with the PDZ-extreme-C-terminus interaction complex. The inset highlights a binding interface similar to that shown in Figure 3A.

Prior research has demonstrated that the extreme-C-terminus of mouse Dvl1 harbours a class III PDZ-binding motif (E/D-X-Φ, where Φ represents hydrophobic residues such as F, I, L, M, or V), whereas in Drosophila it features a class II PDZ-binding motif (Φ-X-Φ) [32]. Previous studies have shown that the mouse Dvl1 adopts an autoinhibited conformation through intramolecular interaction between its extreme-C-terminus and PDZ domain [33]. Class III PDZ binding motifs generally have higher binding affinities than Class II motifs due to a hydrogen bond involving Asp or Glu in Class III, unlike the hydrophobic interface in Class II [51]. Structural analysis of mouse Dvl1 PDZ-extreme-C-terminus complex confirmed this, revealing a hydrogen bond between Asp689 of the extreme-C-terminus and Arg322 of the PDZ domain. Additionally, fluorescence polarization assays indicated that the inhibition constant (K_i_) of Dsh extreme-C-terminus is three times higher than that of mouse Dvl1 [32]. To further explore this, we asses the binding potential of the extreme-C-terminus of Drosophila Dishevelled (Dsh), with the PDZ domains of Dsh and Dvl2. We performed structural alignment of the predicted Dsh PDZ structure with Dvl2 PDZ which yielded a highly significant RMSD of 0.486, indicating strong structural conservation (Figure S6A). However, docking analysis showed that the Dsh extreme-C-terminus does not form a complex with human Dvl2 PDZ and its interaction with Dsh PDZ disrupts the native PDZ conformation, notably causing the loss of beta sheets, which suggests complex instability (Figure S6B).

Our multiple sequence alignment identified a class III PDZ-binding motif (E-F-F-V-D-V) at the extreme-C-terminus of human Dvl2, which is similar to mouse Dvl1 (E-F-F-V-D-I). We performed molecular docking between the Dvl2 PDZ (PDB ID: 3CBZ) domain and its extreme C-terminus to explore the potential for intramolecular interaction. Docking results confirmed that the PDZ domain of Dvl2 can indeed form a stable complex with its extreme-C-terminus (Figure 4B). A 2D interaction map revealed five hydrogen bonds alongside several hydrophobic interactions. The key residues involved in hydrogen bonding were Asp706, Arg718, Gln719, Gly726, and Asn727 from the extreme-C-terminus, and Ile280, Ser281, Gly284, Gln285, and Lys301 from the PDZ domain (Figure S7A). These findings suggest that human Dvl2, like mouse Dvl1, adopts an autoinhibited conformation through intramolecular interaction between its PDZ domain and extreme-C-terminus. From our previous data, we also observed that the extreme-C-terminus Dvl2 interacts with profilin2. This raised an important question, when the extreme-C-terminus is already engaged in an intramolecular interaction with the PDZ domain, can profilin2 still bind to it?

To investigate this, we conducted a molecular docking experiment using the pre-docked PDZ-extreme-C-terminus complex as a single structural unit and docked profilin2 onto it. The results showed that profilin2 could still bind to the extreme-C-terminus. The binding interface included hydrogen bonds between Ser717 and Glu730 of the extreme-C-terminus and Lys28, Asp48, and Lys53 of profilin2 (Figure 4C). Additionally, residues Phe721, Met725, and Phe732, which we previously identified as key to the profilin2-C-terminal Dvl2 (amino acids 536-736) interaction, remained involved in the interaction (Figure S7B). This indicates profilin2 can still interact with the extreme-C-terminus while it remains in an autoinhibited conformation by interacting with the PDZ domain.

To sum up, our data demonstrate that the extreme-C-terminus of human Dvl2 is highly conserved and capable of forming an intramolecular interaction with its PDZ domain, resulting in autoinhibition. Despite this, profilin2 is still able to access and bind the extreme-C-terminus region, highlighting the possibility of a dynamic regulatory mechanism involving competitive or cooperative binding events at this site.

## 3. Discussion

In this study, we identified profilin2 as a novel interacting partner of Dvl2. Our immunofluorescence analysis revealed that Dvl2 and profilin2 are present in both the nucleus and cytoplasm, with predominant co-localization in the perinuclear region (Figure 1A). Additionally, an in vitro pull-down assay verified a direct interaction between endogenous Dvl2 and purified profilin2. Further, immunoblot analysis revealed a Dvl2 band at approximately 90 kDa (Figure S1B). Prior studies have reported that endogenous Dvl2 is distributed throughout HEK293 cells, with a minor fraction in the nucleus, and that Wnt3a treatment enhances its nuclear localization [43]. Also, endogenous Dvl2 maintains a molecular weight of around 90 kDa in untreated cells, while Wnt3a treatment in HEK293T cells increases the molecular weight of Dvl2 to approximately 130 kDa [52]. Given that Wnt3a specifically activates the canonical Wnt pathway [43], these observations raise the possibility of post-translational modifications in canonical signaling. Such mechanisms could, in turn, underlie functional distinctions between canonical and non-canonical Wnt signaling.

We demonstrated that profilin2 interacts with the C-terminal region of Dvl2, and doesn’t interact with PDZ or DEP domain (Figure 2C,D & S2). Intriguingly, our study for the first time reveals that profilin2 exhibits a marked preference for binding to extreme-C-terminus (non-polyproline motif) of Dvl2, even in the presence of a similar GP5 motif (VPPPPPP) (Figure 3B). Previous studies have extensively described Profilin’s interaction with the proline-rich GP5 motif (GPPPPP) of Ena/VASP [47]. Beyond polyproline motifs, Profilin is known to interact with monomeric actin, tubulin, and PIP_2_ [53]. Additionally, profilin interacts with Rho-associated protein kinase 1 (ROCK1), which lacks a polyproline motif, and undergoes phosphorylation at Ser137 [54]. Our findings therefore raise the possibility that profilin engages the C-terminus of Dvl2 through a non-classical interface. Profilin’s preference of polyproline region as a binding surface has been shown with formin and Ena/VASP family proteins [36,55]. It enhances the actin assembly activity of Ena/VASP and formins by allowing the profilin-actin complex to interact with the proline-rich GP5 motif and FH1 domain, respectively. This unexpected selectivity defies the conventional paradigm of profilin-polyproline interactions and suggests the existence of previously unrecognized molecular determinants. An intriguing possibility is that profilin may simultaneously interact with both the FH1 domain of Daam1 and the C-terminal region of Dvl2, thereby regulating actin cytoskeleton with Dvl2-mediated signaling.

The interaction between profilin2 and Dvl2 raises an important question regarding the role of profilin2 in Dvl2-mediated signaling. Profilin1 and profilin2 were shown to share overlapping functions during vertebrate gastrulation. Profilin1 inhibits dorsal blastopore lip closure but does not affect the elongation of Keller’s explants upon depletion [56]. In contrast, depletion of Profilin2 disrupts gastrulation in Xenopus embryo, and inhibit the elongation of Keller’s explants in animal cap assay [40]. This indicates that profilin2 disrupts vertebrate gastrulation by inhibiting PCP pathway. This observation is consistent with our results and indicates that profilin2 and Dvl2 may act along the same regulatory axis within the PCP pathway. Dvl is key regulator of PCP pathway. Three isoforms of Dvl (Dvl1, Dvl2, and Dvl3) are present in mammals. Dvl has been to be involved in Robinow syndrome a rare genetic disorder which causes short limbed dwarfism and genital abnormalities in human [57,58]. Dvl1/2/3 triple mutants are embryonic lethal in mice [59]. In single mutant Dvl1 mice, reduced social interaction is observed, attributed to impaired sensorimotor gating [60]. This effect is likely driven by decreased dendritic arborization, mediated through non-canonical Wnt signalling [61]. Dvl2 mutant mice exhibit cardiac defects and neural tube closure abnormalities, characteristic of disrupted non-canonical Wnt signalling [62]. Dvl1/2 double mutant mice display more severe cardiac abnormalities and neural tube defects compared to single Dvl2 mutant mice [59]. Together, these findings highlight the essential contributions of both profilin2 and Dvl2 to the regulation of non-canonical Wnt signalling. This raises the possibility that a direct interaction between profilin2 and Dvl2 could represent a key mechanism for fine-tuning non-canonical Wnt pathway activity.

Our multiple sequence alignment of the human Dvl2 extreme-C-terminus revealed striking evolutionary conservation with mouse (Dvl1, Dvl2, Dvl3), Xenopus (Dvl1, Dvl2, Dvl3), and zebrafish (Dvl1, Dvl2, Dvl3), and retains over 60% similarity even with more distant species such as ascidian (Dvl) and Hydra (Dsh) (Figure 4A). Interestingly, this high level of conservation does not extend to Drosophila. We further demonstrate that Drosophila Dsh PDZ domain forms an unstable complex with its own extreme-C-terminus (Figure S6B). Moreover, the extreme-C-terminus of Dsh fails to interact with the PDZ domain of human Dvl2. These results suggest that Dsh is unlikely to adopt an autoinhibited conformation through PDZ-extreme-C-terminus interactions, raising the possibility that alternative regulatory mechanisms govern Dsh activity. Overexpression of a Dsh-GFP construct in Drosophila larvae induces Wg (Wnt homolog)-independent membrane localization of Dsh in wings [63]. This indicates involvement of Dsh in PCP without the effect of Wnt ligands. Future studies will be required to clarify the alternative mechanisms regulating Dsh and to determine whether PCP signaling in Drosophila fundamentally diverges from that in vertebrates.

We further revealed human Dvl2 adopts an autoinhibited conformation via interaction between its PDZ and extreme-C-terminus (Figure 4B). Notably, profilin2 was associated with the autoinhibited extreme-C-terminus-PDZ complex (Figure 4C). A similar mechanism has been described for mouse Dvl1, where the extreme C-terminus forms an intramolecular complex with its PDZ domain, thereby establishing an autoinhibited state [32]. This parallel supports our finding that human Dvl2 can also adopt such an autoinhibitory configuration. Functional relevance of the C-terminal region has been demonstrated in zebrafish embryos. When the truncated construct lacking the extreme-C-terminus and only C-terminus was overexpressed in zebrafish embryos, it increases membrane localization of Dvl1, a hallmark of PCP pathway activation, and causes significant shortening of the anterior-posterior (A-P) axis in zebrafish embryos, indicating impaired PCP signalling [33]. These findings highlight the critical importance of the extreme-C-terminus in maintaining Dvl2’s autoinhibited conformation and suggest that disrupting this regulatory mechanism can profoundly alter PCP signaling outcomes. In the autoinhibited state, Dvl mainly exposes its DIX domain, favoring canonical Wnt signalling, whereas deletion of the extreme-C-terminus may release this restraint and allow PDZ, DEP, and C-terminal regions to engage downstream effectors of the non-canonical pathway. This raises intriguing possibilities, is the extreme C-terminus acting as a molecular switch that toggles between canonical and non-canonical Wnt outputs? How profilin2 by associating to the autoinhibited complex, fine-tune this switch to regulate Dvl2 functionality? How exactly the extreme-C-terminus facilitates non-canonical signalling, and the extent to which profilin2 contributes to this regulation, remain open questions that warrant further investigation.

Our research reveals that profilin2 interacts with the extreme-C-terminus of Dvl2, which regulates the PCP pathway. Studies show that profilin1 colocalizes with Daam1 at actin stress fibers in response to Wnt stimulation [56]. Depletion of profilin1 in NIH3T3 cells inhibits Wnt and Daam1-mediated stress fiber formation and disrupts gastrulation in Xenopus embryos by impairing the PCP pathway [56]. This indicates that profilin1 acts downstream of Daam1 in this pathway. In contrast, silencing profilin1 in T24M bladder cancer cell line suppresses Wnt/Ca^2+^ signalling by inhibiting PLC [41]. In the Wnt/Ca^2+^ pathway, Dvl2 promotes cytoskeletal reorganization by releasing intracellular Ca^2+^ ions via trimeric G-protein and PLC, independent of Daam1 [29]. Another study has shown depletion of Profilin2 disrupts gastrulation by inhibiting PCP pathway [40]. Collectively, these findings suggest that profilin is involved in both branches of non-canonical Wnt signalling (PCP and Wnt/Ca^2+^ pathways). Based on our findings, we hypothesize that profilin2 may operates upstream of Daam1 along the Dvl2 axis. This creates ambiguity regarding the precise position of profilin in non-canonical Wnt signalling (Figure 5). This also points to non-redundant functions of profilin isoforms in non-canonical Wnt signalling. Further investigation is needed to clarify how profilin isoforms regulate these pathways in a Dvl-dependent or -independent manner. Previous studies have shown that profilin interacts with Daam1 and enhances its actin filament elongation activity [56]. Dvl2, in turn, activates Daam1 [64]. In this study, we demonstrate that profilin2 also interacts with Dvl2. These findings suggest two possible regulatory scenarios. In the first, profilin2 engages with Dvl2, leading to its activation. The activated Dvl2 then triggers Daam1 activation, after which profilin2 may also interact with Daam1 to further modulate non-canonical Wnt signalling (Figure 5). In the second, profilin2 could simultaneously associate with both Dvl2 and Daam1, forming a trimeric complex that fine-tunes pathway activity (Figure 5). Together, these observations point to a multilayered regulatory framework in which Dvl2, Daam1, and profilin2 participate in both sequential and cooperative interactions, ensuring precise control of PCP signaling.

**Figure. 5.**
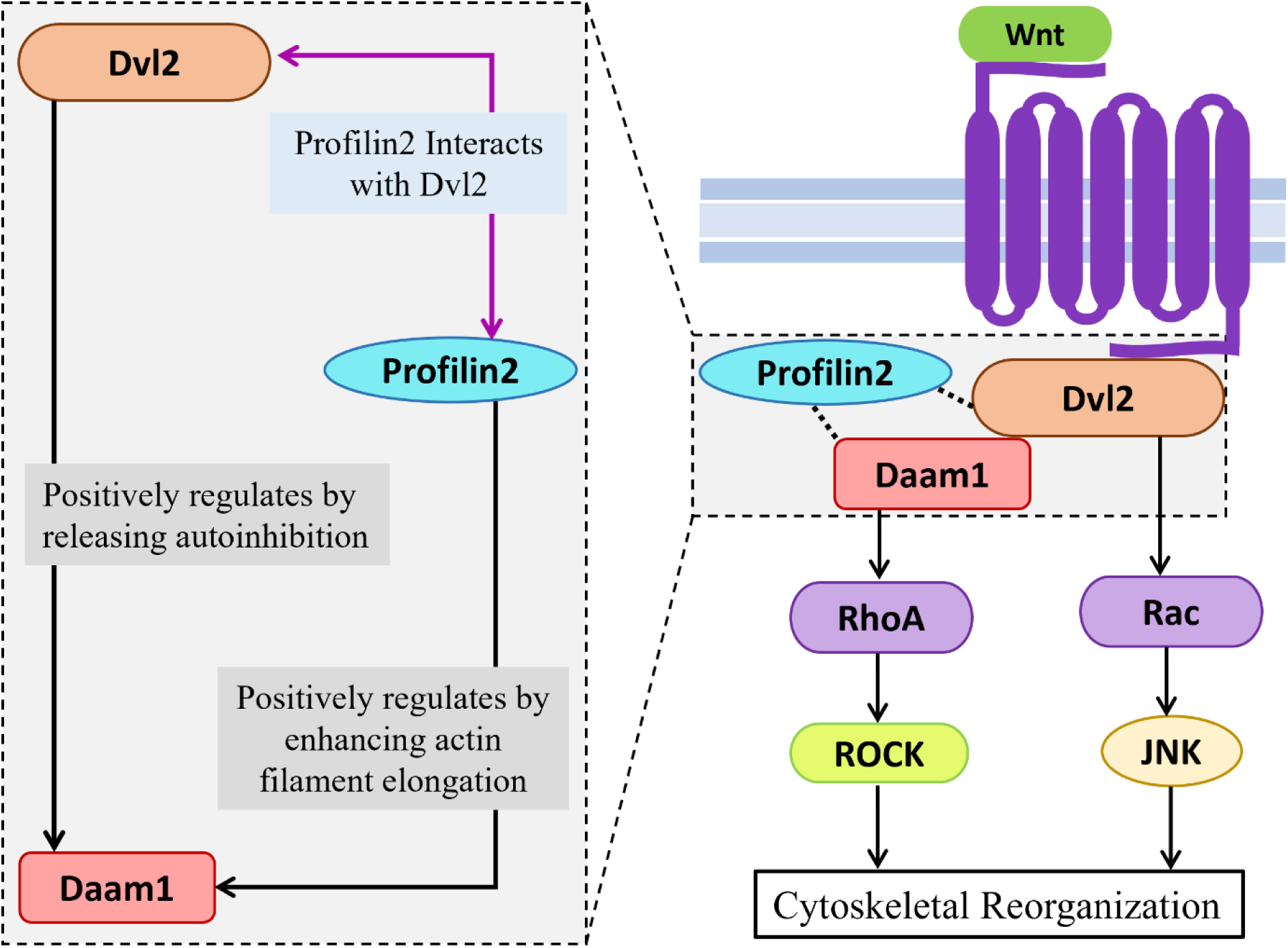
Positional ambiguity of Profilin2 in non-canonical Wnt signalling. The diagram on the right illustrates non-canonical Wnt signaling, where Wnt binds to the Frizzled (Fz) receptor, prompting Dvl2 to form a complex with Daam1 and RhoA. This interaction activates RhoA, which subsequently triggers Rho-associated kinase (ROCK), leading to cytoskeletal reorganization. Profilin2 is known to interact with Daam1, and here we show that it also interacts with Dvl2. This raises questions about profilin2’s precise role and positional ambiguity in non-canonical Wnt signaling, as emphasized in the inset on the right.

## 4. Materials and method

### 4.1. Plasmid construct and cloning

The human Dvl2 was obtained from pCMV-SPORT6-Dvl2 plasmid (Clone ID #3852554, UniProtKB ID O14641; Organism-*Homo sapiens*). We subsequently generated three constructs of human Dvl2, Dvl2 (amino acids 182-389), Dvl2 (amino acids 390-536), and the C-terminal Dvl2 (amino acids 536-736) (Figure 2A), all subcloned into 6x-His tag containing pET28a (+) vector (Novagen). Additionally, we created a polyproline region-deleted construct, ΔPP C-terminal Dvl2 (amino acids 536-736) (Figure 3B), using the C-terminal Dvl2 (amino acids 536-736) fragment as a template. Full-length mouse Profilin2 (PFN2; PDB-ID 2V8C; Organism-<u>Mus musculus</u>) was subcloned into the pGEX-4T-3 vector, as previously described [65].

### 4.2. Cells and Antibodies

HEK293T cells (CRL-3216, Lot Number 70049877), obtained from the American Type Culture Collection, were cultured in Dulbecco’s Modified Eagle Medium (DMEM) containing 2 mM L-glutamine, 1% penicillin/streptomycin, and 10% fetal bovine serum (FBS).

Antisera against Dvl2 and Profilin2 were generated by immunizing BALB/c mice and New Zealand White rabbits with purified His-tagged Dvl2 (amino acids 536-736) and GST-tagged profilin2, respectively. The immunization protocol, lasting 70 days, was approved by the Institutional Animal Ethics Committee (IAEC) under protocol number IISERK/IAEC/2022/024 for His-tag Dvl2 (amino acids 536-736) and IISERK/IAEC/AP/2022/85 for GST-tag profilin2, respectively. Terminal bleeds were validated using western blot analysis with recombinant proteins and cell lysates (Figure S1).

### 4.3. Immunofluorescence staining

HEK293T cells (5 × 10⁴ cells/mL) were directly seeded onto poly-L-lysine-coated coverslips and incubated to allow attachment. Cells were fixed and permeabilized post-attachment using a 1:1 acetone-methanol mixture for 15 minutes at −20 °C. Blocking was performed with 2% bovine serum albumin (BSA) in PBS for 2 hours. Primary antibody incubation was carried out in 1% BSA-PBST (PBS with 0.075% v/v Tween-20) for 2 hours at room temperature, followed by incubation with fluorophore-tagged secondary antibodies for 1 hour at room temperature. Nuclei were stained with 4’,6-diamidino-2-phenylindole (DAPI) included in the fluoroshield mounting solution. After each step, cells were washed with PBS. The combination of anti-Dvl2 antisera (1:200) plus Alexa Fluor 488-conjugated anti-mouse IgG (1:1000) was used to visualize localization of endogenous Dvl2. While anti-profilin2 antisera (1:500) plus Alexa Fluor 568-conjugated anti-rabbit IgG (1:1000) combination was used to visualize localization of endogenous Profilin2. Leica SP8 confocal microscope system fitted with a 63×/1.40 N.A. oil immersion objective (HC PL APO CS2 63×/1.40 OIL) was used to capture the image.

### 4.4. Protein expression and purification

Constructs of Dvl2 containing PDZ and DEP domains-Dvl2 (amino acids 182-389) and Dvl2 (amino acids 390-536) were transformed into BL21(DE3) *E. coli*, while C-terminal Dvl2 (amino acids 536-736) and ΔPP C-terminal Dvl2 (amino acids 536-736) were transformed into BL21(DE3)RP *E. coli*. The cells were cultured in LB medium at 37°C with 30 µg/mL kanamycin until reaching an OD of 0.5. Protein expression was induced with 0.5 mM IPTG at 18°C for 12 hours. Cells were then harvested via centrifugation. All His and GST-tag proteins are purified following the previously published protocol with slight modifications [66].

#### 4.4.1. His-tag protein purification

All Dvl2 fragments, tagged with 6x-His, were resuspended in His-lysis buffer (50 mM Tris-Cl pH 8, 100 mM NaCl, 30 mM imidazole pH 8, 5 mM MgCl_2_, 0.25% IGEPAL, 1X PIC [phenylmethylsulfonyl fluoride, benzamidine hydrochloride, leupeptin, aprotinin, pepstatin A], and 0.5 mM DTT) and sonicated with 5% amplitude for 4 minutes with 20 seconds on 60 seconds off cycle. The lysates were centrifuged at 14,000 rpm for 10 minutes. Supernatants were incubated with Ni-NTA beads for 2 hours with 5 rpm rotation. Beads were washed with His-wash buffer (50 mM Tris-Cl pH 8, 300 mM NaCl, 30 mM imidazole pH 8, 5 mM MgCl_2_), and proteins were eluted using His-elution buffer (50 mM Tris-Cl pH 8, 20 mM NaCl, 350 mM imidazole pH 8, 5% glycerol).

#### 4.4.2. GST-tag protein purification

For profilin2 expression, the profilin2 construct was transformed into BL21(DE3) cells and grown at 37°C in LB medium with 100 µg/mL ampicillin until an OD of 0.5. Expression was induced with 0.5 mM IPTG at 25°C for 8 hours. Cells were harvested and resuspended in GST lysis buffer (50 mM Tris-Cl pH 7.5, 100 mM NaCl, 1 mM EDTA, 5 mM MgCl_2_, 0.25% IGEPAL, 1X PIC, 2 mM DTT), sonicated with 1% amplitude for 2 minutes with 10 seconds on 30 seconds off cycle, and centrifuged at 14,000 rpm for 10 minutes. The supernatant was incubated with beads, which were washed with GST wash buffer (50 mM Tris-Cl pH 7.5, 300 mM NaCl, 1 mM EDTA, 5 mM MgCl_2_) and eluted with GST elution buffer (50 mM Tris-Cl pH 7.5, 100 mM NaCl, 10 mM glutathione, 5% glycerol).

For general purpose (like ELISA and antibody development), dialysis was performed in HEKG_5_ buffer (20 mM HEPES, 1 mM EGTA, 50 mM KCl, 5% glycerol) for 4 hours. For SPR analysis, all proteins were dialyzed in HBS-N buffer (10 mM HEPES pH 7.4, 150 mM NaCl) (Cytiva) for 4 hours. All protein purification steps were conducted on ice or at 4°C.

### 4.5. GST Pulldown assay

To investigate the direct interaction between Dvl2 and profilin2, we performed a GST pull-down assay using purified GST-tagged profilin2 as the bait and HEK293T cell lysate as the source of the prey protein (Dvl2). HEK293T cells were lysed in lysis buffer (20 mM Tris pH 7.5, 150 mM NaCl, 1 mM EDTA, 0.5% IGEPAL, and 1 mM PMSF) by passing the suspension through a 31-gauge needle 60 times. Following lysis, the cell suspension was centrifuged at 12,000 rpm for 15 minutes at 4°C to remove cell debris. The resulting supernatant was then incubated with a 50% slurry of glutathione-agarose beads for 2 hours at 4°C to pre-clear the lysate. This step helps eliminate proteins that may non-specifically bind to the beads. After pelleting the beads, the pre-cleared lysate was divided into two aliquots and incubated overnight at 4°C with 25 µg of GST alone and GST-tagged profilin2.

To capture the bait-prey (Profilin2-Dvl2) complexes, glutathione-agarose beads were added to each reaction and incubated for 2 hours. The beads were collected by centrifugation and washed three times with ice-cold lysis buffer to remove non-specifically bound proteins. All steps were carried out on ice or at 4°C. Bound proteins were eluted by boiling the beads in Laemmli buffer and analysed by western blotting. Proteins were resolved on a 10% SDS-PAGE gel and transferred onto a nitrocellulose membrane using transfer buffer (25 mM Tris pH 7.5, 192 mM glycine, 0.2% SDS, 10% methanol) under a constant electric field of 12V. The bait-prey complex was detected by incubating the blot with mouse anti-Dvl2 antibody (1:500 dilution) plus HRP-conjugated goat anti-mouse IgG secondary antibody (Invitrogen, 1:50,000 dilution) combination.

### 4.6. Enzyme-linked immunosorbent assay (ELISA)

To identify the region of Dvl2 that interacts with profilin2, an ELISA-based assay was conducted using previously published protocol [46]. The purified Dvl2 (amino acids 182-389), Dvl2 (amino acids 390-536), and Dvl2 (amino acids 536-736) fragments were individually coated onto ELISA plates (Maxisorp surface) at a concentration of 10 µg per well, with PBS serving as a negative control. Plates were incubated overnight at 4°C. All subsequent steps were performed at room temperature. The wells were incubated with 5% BSA for 2 hours, followed by incubation of purified profilin2 in a concentration-dependent manner for 2 hours at room temperature. Detection of the Dvl2–Profilin2 complex involved three steps: (1) addition of a rabbit anti-profilin2 antibody (1:1000 dilution) for 2 hours, (2) incubation with an HRP-conjugated goat anti-rabbit IgG secondary antibody (Invitrogen, 1:10000 dilution) for 45 minutes, and (3) development of color using Tetramethylbenzidine (TMB; Sigma Aldrich, 1X dilution) for 15 minutes. The reaction was stopped by adding 5N H₂SO₄. Between each step (up to the TMB addition), wells were washed with PBST (0.02% Tween-20 in 1X PBS). Absorbance at 450 nm was measured using a microplate reader (Epoch2), and an absorbance vs. concentration (µg/mL) curve was generated using GraphPad Prism 8.

As initial results indicated that the C-terminal region of Dvl2 (amino acids 536-736) mediates the interaction with Profilin2, a reverse ELISA was performed to validate this finding using the same protocol. In this experiment, wells were coated with profilin2 (10 µg per well), and purified Dvl2 (amino acids 536-736) was added as the analyte in a concentration-dependent manner. Detection of the profilin2-Dvl2 complex involved incubation with a mouse anti-Dvl2 antibody (1:800 dilution), followed by an HRP-conjugated goat anti-mouse IgG secondary antibody (Invitrogen, 1:50000 dilution). As before, TMB was used for color development, and absorbance at 450 nm was recorded and plotted using GraphPad Prism 8.

### 4.7. Surface plasmon resonance (SPR)

The binding kinetics of Dvl2 (amino acids 536-736) and ΔPP C-terminal Dvl2 (amino acids 536-736) with profilin2 were evaluated using amine-coupling surface plasmon resonance (SPR) on a Biacore T200 (Cytiva) with slight modifications from previously published protocol [49]. The CM5 sensor chip (Series S) surface was treated with an EDC/NHS mixture which will activate the surface by creating reactive succinimide ester groups. Purified Profilin2 (25 µg/mL) in sodium acetate buffer (pH 5) was immobilized in the presence of running buffer HBS-EP (0.01 M HEPES, 0.15 M NaCl, 0.03 M EDTA, 0.05% surfactant P20, pH 7.4) at a flow rate of 30 µL/min. Residual active succinimide ester groups were blocked with ethanolamine, and a non-immobilized reference cell was used for blank correction. Purified Dvl2 (amino acids 536-736) and ΔPP C-terminal Dvl2 (amino acids 536-736) were passed over the sensor surface in a concentration-dependent manner. HBS-EP buffer served as the running buffer, with 10 mM glycine (pH 2.5) used for regeneration after each cycle. The association phase lasted 180 s or 120 s, with a dissociation phase of 180 s. Experiments were conducted at 25°C, with the sample compartment maintained at 18°C. Sensograms were analyzed using Biacore Evaluation Software (version 2.0), and fitted to a 1:1 Langmuir binding model to determine equilibrium dissociation constants (K_D_), association rates (k_a_), and dissociation rates (k_d_).

### 4.8. Molecular docking

#### 4.8.1. Complex of C-terminal Dvl2 and Profilin2

To identify the region of Dvl2 involved in Profilin2 binding, molecular docking studies were carried out. A 3D structure of the C-terminal Dvl2 (amino acids 536-736) (UniProt ID: O14641) was predicted using AlphaFold Colab. The best model was subsequently docked with Profilin2 (PDB ID: 2V8C) using the HDOCK webserver (http://hdock.phys.hust.edu.cn/).

#### 4.8.2. Complex of Dvl2 PDZ Domain and C-terminal Dvl2

To explore the potential autoinhibitory interaction between the PDZ domain and the extreme C-terminal region of Dvl2, molecular docking was performed using the Dvl2 PDZ domain (PDB ID: 3CBZ) and the previously modeled C-terminal Dvl2 (amino acids 536-736). Docking was carried out using the HDOCK webserver.

#### 4.8.3. Ternary Complex of PDZ Domain, C-terminal Dvl2, and Profilin2

To assess whether Profilin can still bind to the autoinhibitory Dvl2 complex, a ternary docking study was conducted. The pre-docked complex of the Dvl2 PDZ domain and C-terminus Dvl2 (amino acids 536-736) was treated as a single entity, to which Profilin2 was docked using the HDOCK webserver. All resulting protein complexes and their interaction interfaces were visualized using PyMOL (v2.5) (https://pymol.org/). Furthermore, 2D interaction maps were generated with LigPlot+ (https://www.ebi.ac.uk/thornton-srv/software/LigPlus/), highlighting key molecular interactions such as hydrogen bonds, hydrophobic contacts, and ionic interactions.

## Data availability

All datasets that support the results of this study are available from the corresponding author upon reasonable request.

## Acknowledgements

Saikat and Shubham extend heartfelt thanks to the Council of Scientific & Industrial Research and University Grants Commission for their fellowships. SM acknowledges DST-FIST support for the SPR facility at the Central Analytical Instrumentation Facility. We would like acknowledge Mr. Amaresh Jana and Mr. Sukanta Murmu for their critical comments on the manuscript.

## Author contributions

Saikat and Shubham contributed equally to this work. Saikat and Shubham: Data curation; investigation; visualization; validation; methodology; writing-original draft; writing-review and editing. S.M.: Conceptualization; funding acquisition; project administration; writing-original draft; writing-review and editing.

## Supporting Information

**Table S1:** Strains and plasmids used for this study.

| **Strains** | **Source** |
| --- | --- |
| XL10 Gold | #200314, Stratagene, Agilent Technologies |
| BL21(DE3) | #200131, Agilent Technologies |
| BL21(DE3)-RP | #230250 Agilent Technologies |
| **Plasmids** | **Source** |
| pET28+ | #69864, Novagen, Merck Life Science |
| pGEX-4T3 | # V010916, GE Healthcare Life Sciences |
| GST-Profilin2 | P. Dutta et al., Experimental Cell Research,<br>2017[65] |

**Table S2:**
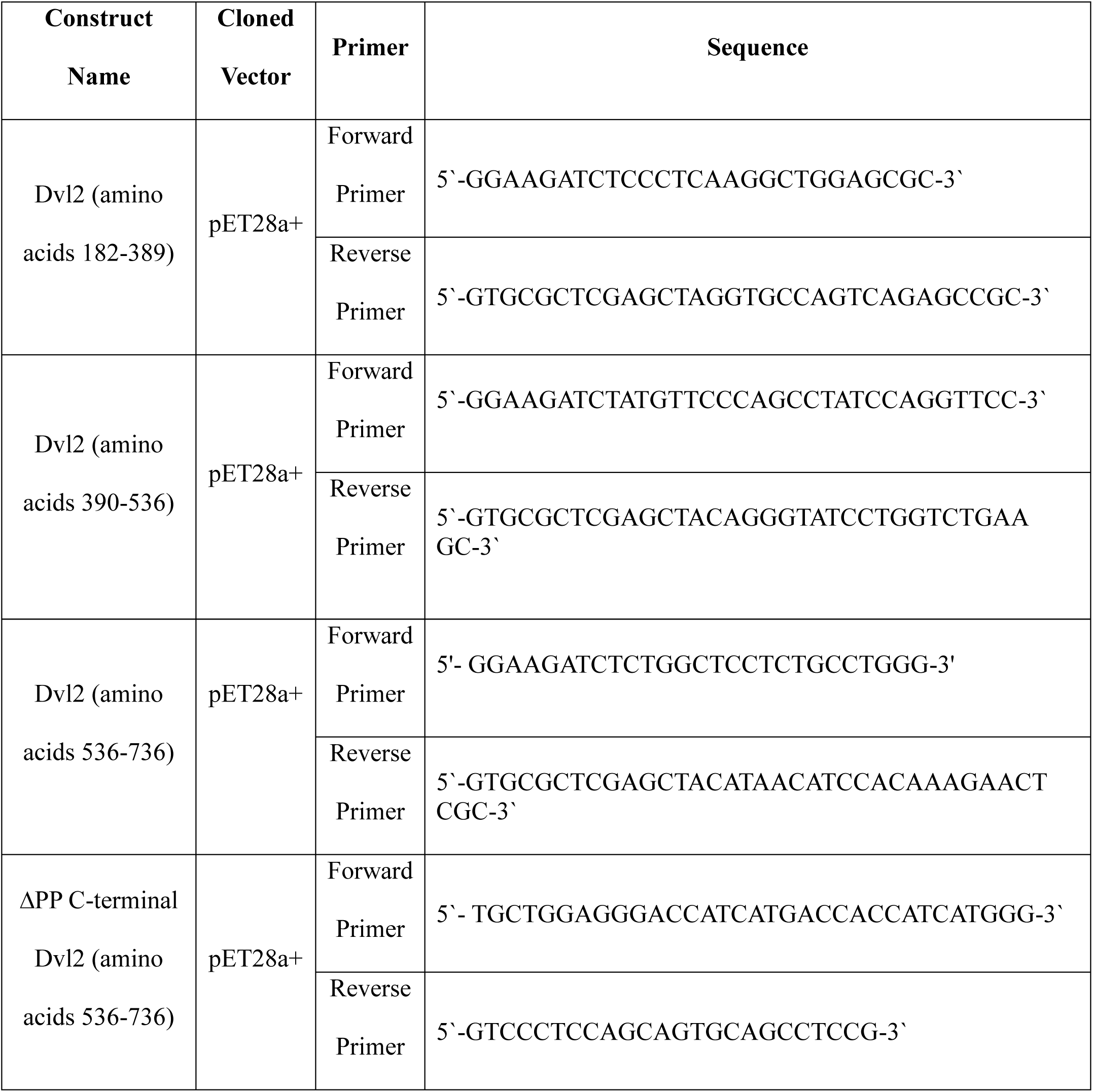
Oligonucleotides list of all the constructs used for this study.

**Figure S1.**
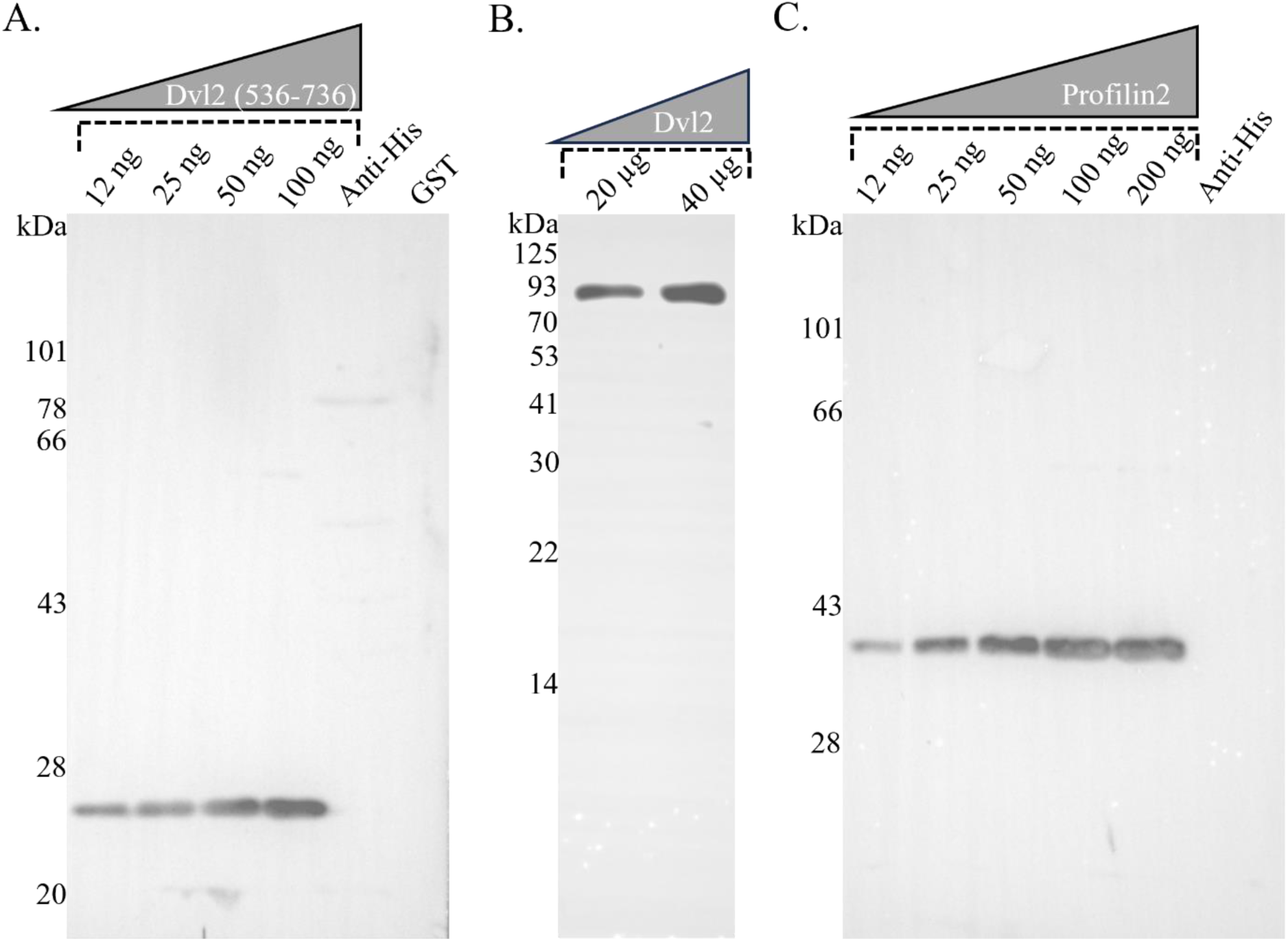
Western blot analysis for sensitivity testing of Dvl2 and Profilin2 antibodies. (A) The sensitivity of the mouse-raised anti-Dvl2 antibody (1:800) was evaluated by western blot using four different concentrations of purified Dvl2 (amino acids 536-36), along with Anti-His-tag protein and GST controls. (B) Additionally, 20 µg and 40 µg of HEK293T cell lysate were used to assess endogenous Dvl2 expression in HEK293T cells. (C) The sensitivity of the rabbit-raised anti-profilin2 antibody (1:1000) was tested using five different concentrations of purified profilin2, with His-tag protein control included.

**Figure S2.**
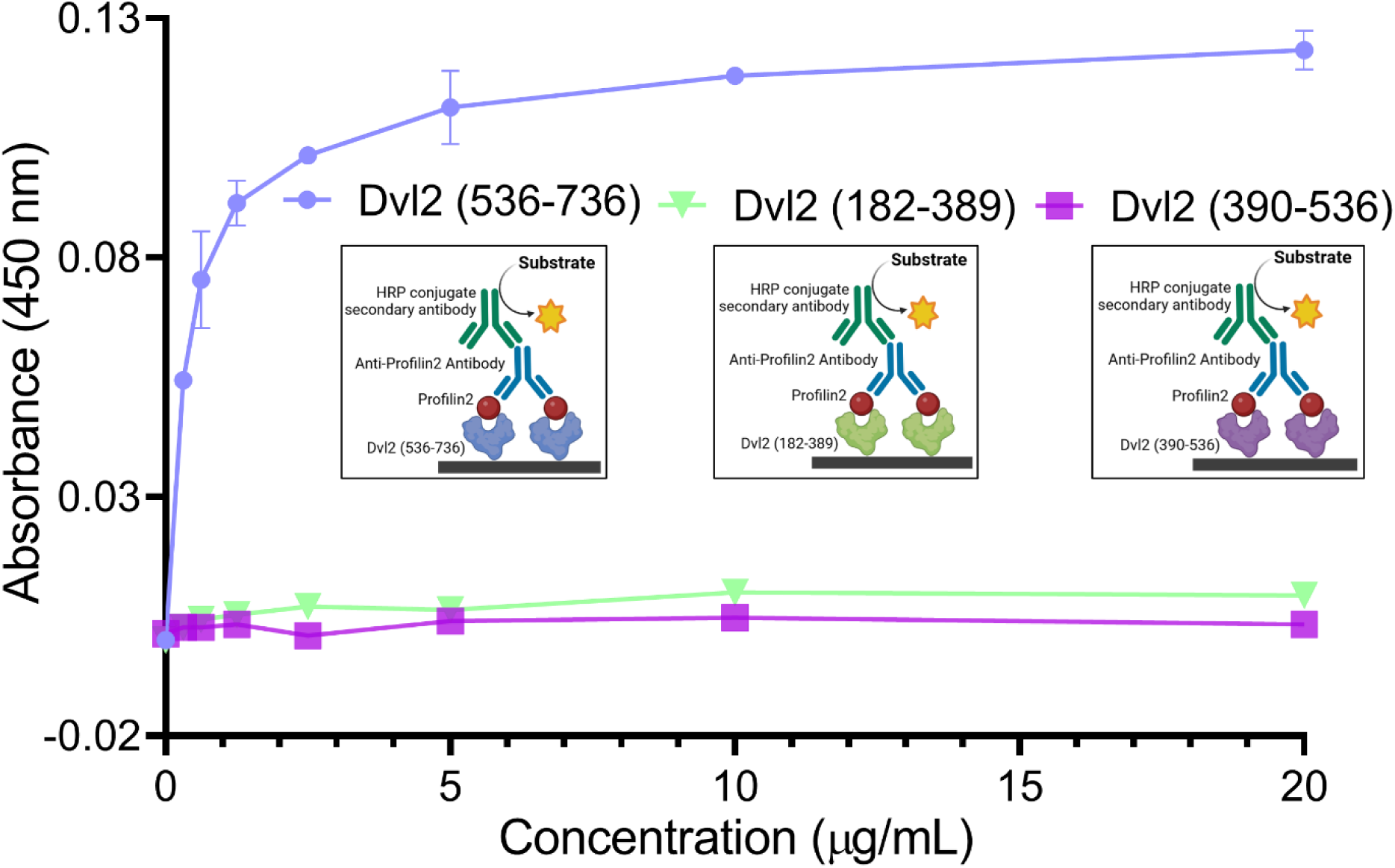
Profilin2 and Dvl2 interaction is mediated by the C-terminal region of Dvl2, excluding the PDZ and DEP domains. ELISA analysis of Dvl2 fragments and Profilin2 interaction. Purified 6x-His-tagged Dvl2 fragments-(amino acids 182-389), (amino acids 390-536), and (amino acids 536-736) were immobilized on ELISA plates. Profilin2 was used as the analyte for each experimental set up. Following incubation, anti-profilin2 antibody (1:1000) was added, followed by HRP-conjugated secondary anti-IgG antibody. TMB substrate was used for color development, and absorbance was measured at 450 nm. Absorbance values were plotted on the Y-axis, and analyte concentrations were plotted on the X-axis.

**Figure S3.**
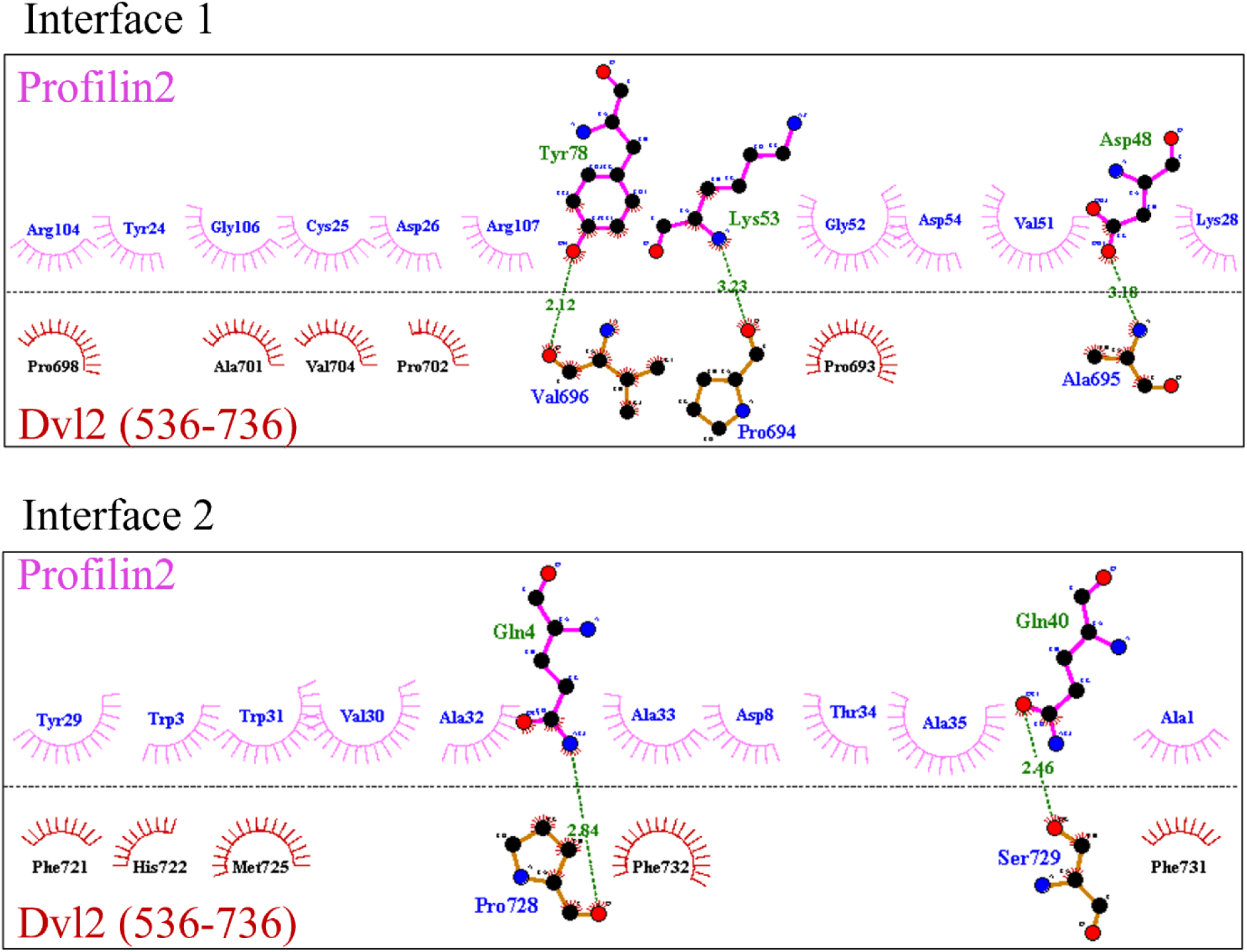
2D interaction map of the Profilin2-Dvl2 (amino acids 536–736) complex. The map illustrates the binding interface, highlighting amino acid residues involved in hydrogen bonding (indicated by green dotted lines) and hydrophobic interactions (represented by red and pink spheres). Two distinct interaction interfaces are shown.

**Figure S4.**
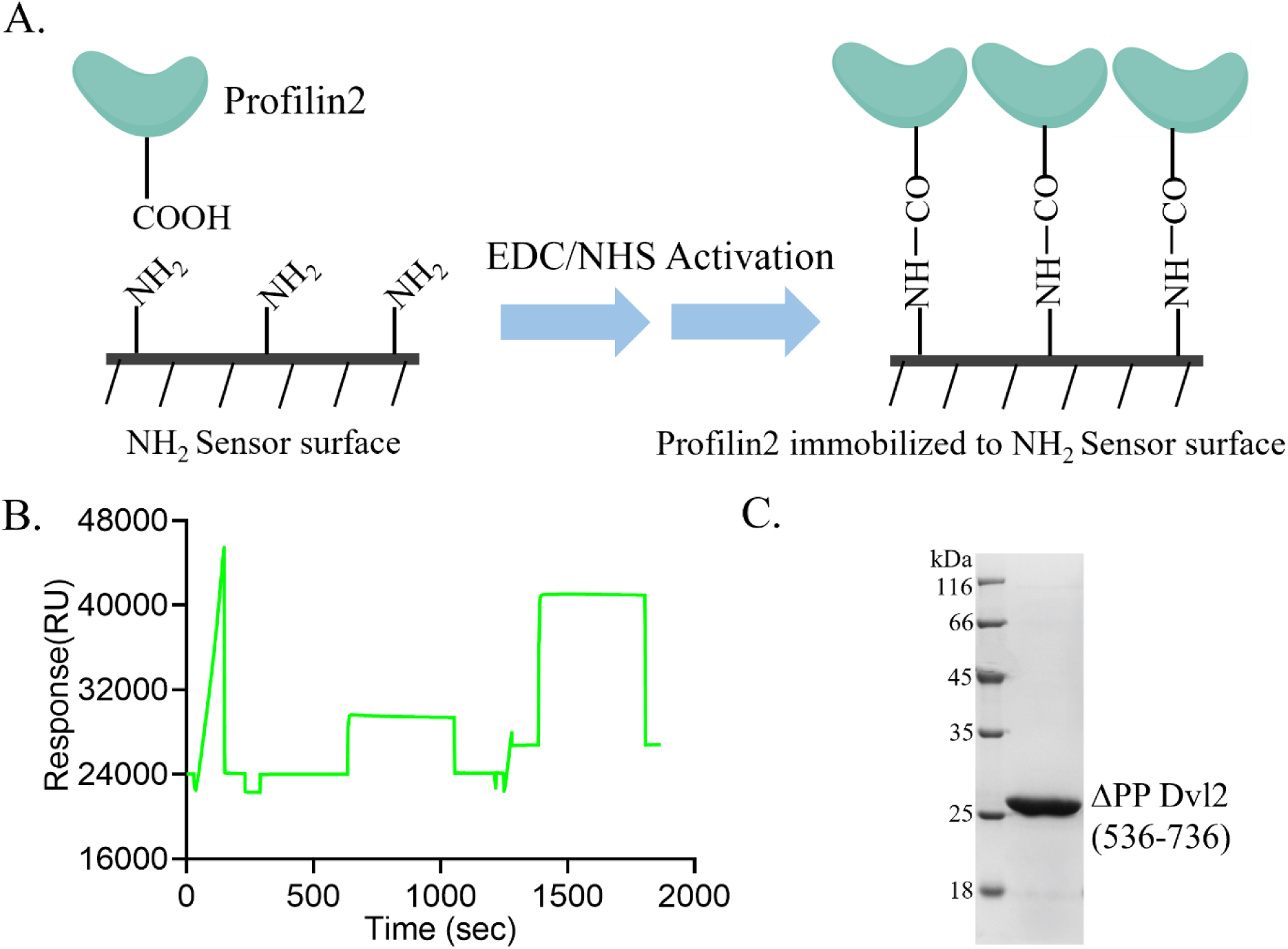
Profilin2 immobilization and ΔPP C-terminal Dvl2 purification for SPR analysis. (A) Schematic representation of amine-coupling-based immobilization of Profilin2 in SPR. (B) Immobilization of profilin2 on reference channel 2. The sensor chip surface was activated using EDC/NHS chemistry, followed by application of profilin2 (25 μg/mL in sodium acetate buffer, pH 5.0) for covalent attachment. Surface deactivation was performed using ethanolamine. (C) Coomassie-stained 10% SDS-PAGE showing purified ΔPP C-terminal Dvl2 (amino acids 536-736) fragment.

**Figure S5.**
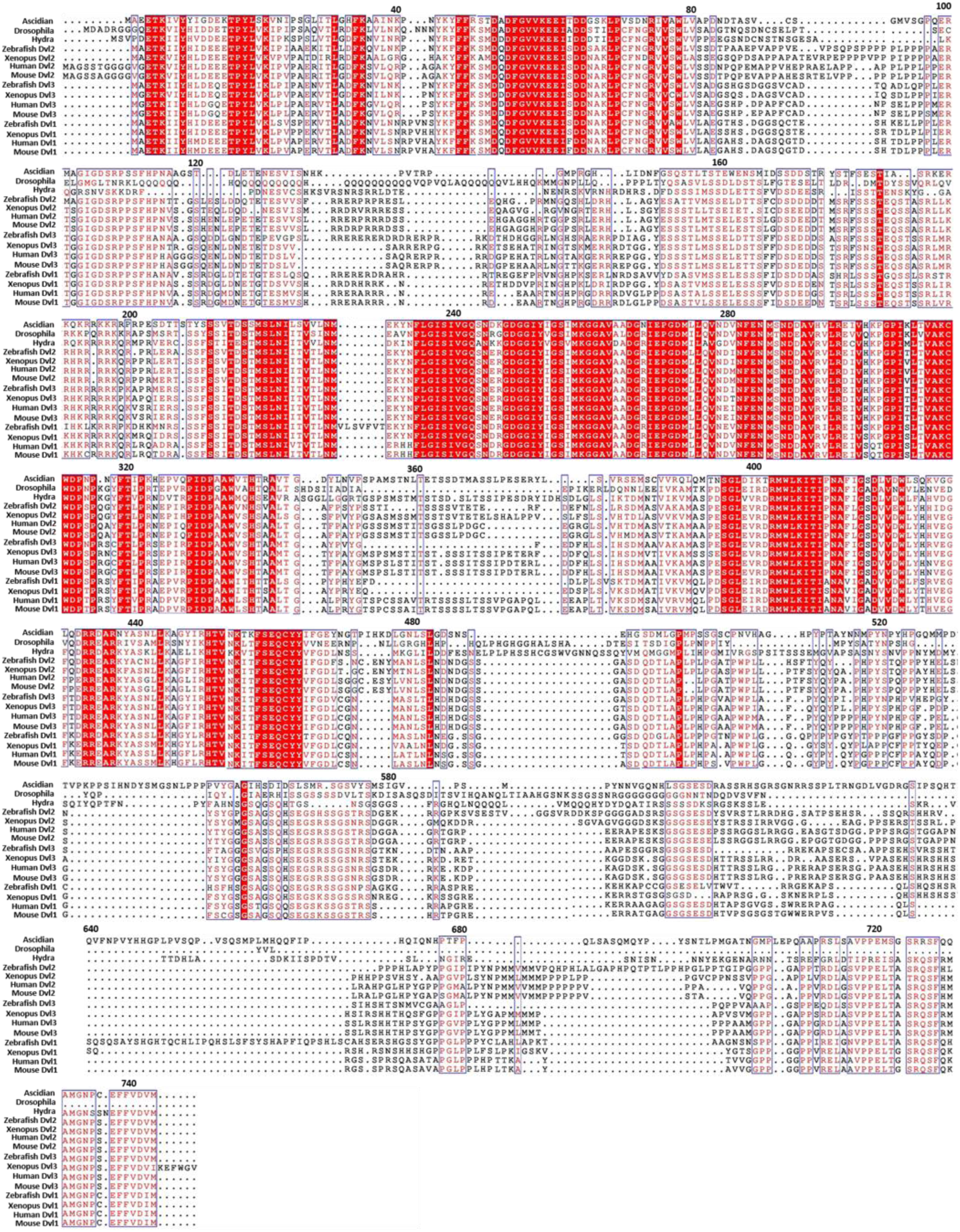
Multiple sequence alignment of Dishevelled proteins from Hydra, ascidian, zebrafish, Drosophila, Xenopus, mouse, and human. Fully conserved residues are highlighted as white letters within red boxes, while residues conserved within specific animal classes are shown as red letters within blue boxes.

**Figure S6.**
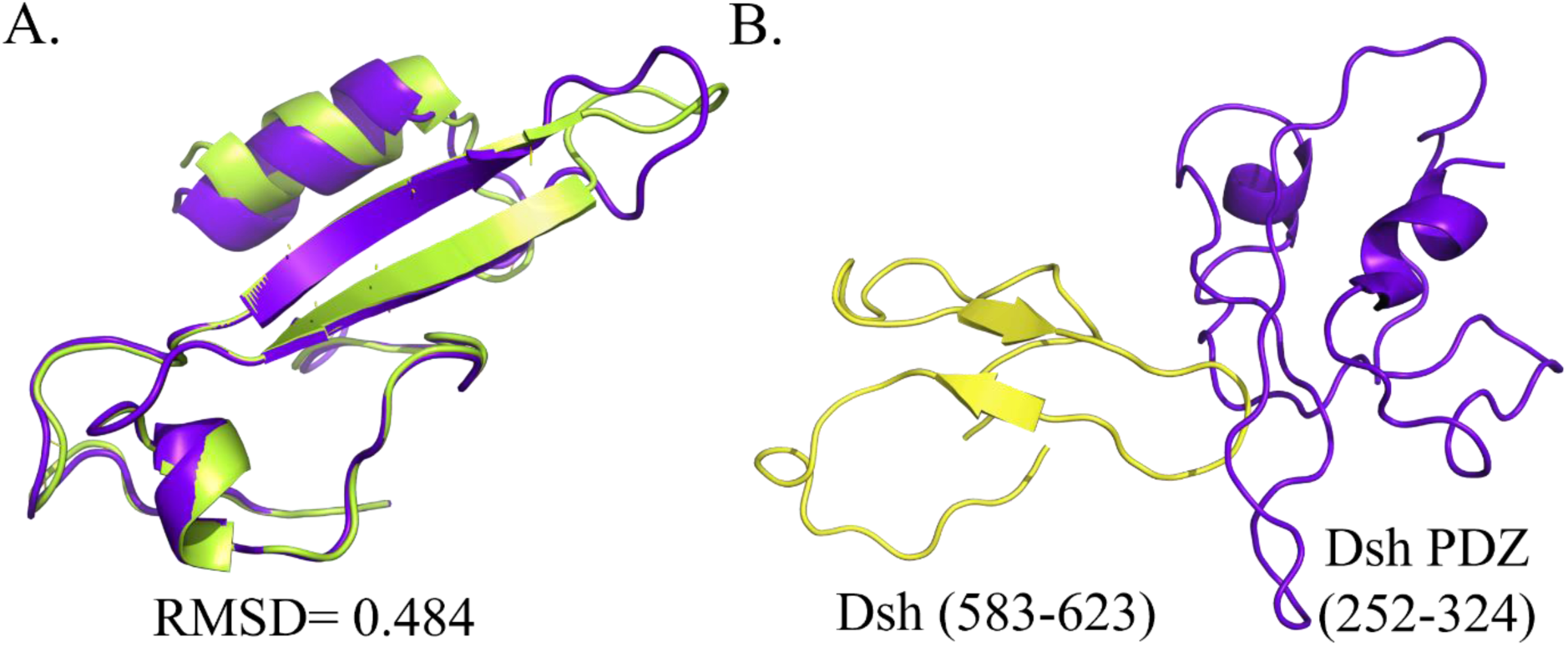
No intramolecular interaction detected between the Drosophila PDZ domain and its extreme C-terminus. (A) Superimposition of PDZ domains from human Dvl2 (in lime green) and Drosophila Dsh (in violetpurple) reveals strong structural conservation. (B) The top-ranked docked complex of the Drosophila PDZ domain (in violetpurple) with its extreme-C-terminus (in yellow) indicates an unstable interaction.

**Figure S7.**
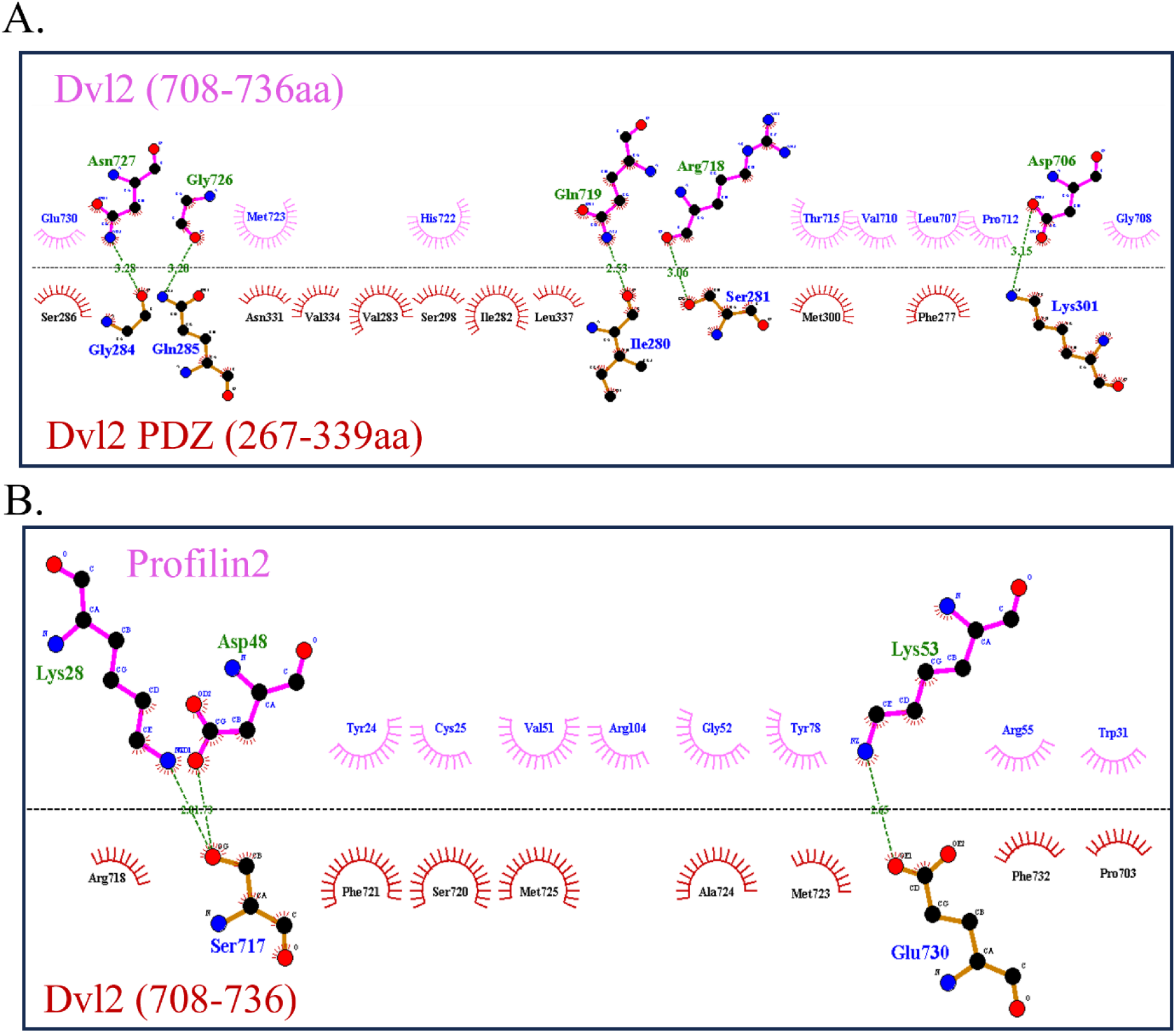
LigPlot-generated 2D interaction maps of the PDZ–extreme C-terminus– Profilin complex. (A) PDZ–extreme C-terminus complex and (B) extreme C-terminus– Profilin complex. Amino acid residues involved in hydrogen bonding are shown as sticks, with hydrogen bonds indicated by green dashed lines, while hydrophobic interactions are represented by red and pink circles.

